# Life history parameters in acellular extrinsic fiber cementum microstructure

**DOI:** 10.1101/528760

**Authors:** Marija Edinborough, Sarah Fearn, Matthew Pilgrim, Andrijana Cvetković, Branko Mihailović, Rade Grbić, Kevan Edinborough

**Affiliations:** Melbourne Dental School, Melbourne Dental School, The University of Melbourne, Melbourne, Victoria, Australia; Royal School of Mines, Department of Materials, Imperial College London, London, United Kingdom; Eastman Dental Institute, University College London, London, United Kingdom; Institute of Ophthalmology, University College London, London, United Kingdom; Faculty of Medicine, University of Priština, Kosovska Mitrovica, Serbia; Department of Anthropology, University of British Columbia, Vancouver, Canada

**Author notes:** These authors contributed equally to this work (Marija Edinborough, Kevan Edinborough). These authors also contributed equally to this work (Sarah Fearn, Matthew Pilgrim).

## Abstract

Life-history parameters such as pregnancies, skeletal trauma, and renal disease have previously been identified from hypomineralized growth layers (incremental lines) of acellular extrinsic fiber cementum (AEFC). The precise periodicity of these growth layers remains vaguely approximated, so causal life-history explanations using tooth cementum cannot yet be rigorously calculated or tested. On the other hand, we show how life history parameters in AEFC can be identified by two contrasting elemental detection methods. Based on our results we reject the possibility of accurate estimation of pregnancies and other life history parameters from cementum using scanning electron microscopy alone. Here, we propose a new methodological approach for cementum research, Time-of-Flight Secondary Ion Mass Spectrometry (ToF-SIMS), to measure degree and distribution of mineralization of cementum growth layers. Our results show that Tof-SIMS can significantly increase our knowledge of cementum composition and is therefore a powerful new tool for life history researchers.

## Introduction

Acellular extrinsic fiber cementum (AEFC) is deposited in a regular annual rhythm in the form of incremental lines around the roots of human teeth, with varying degrees of mineralization [1, 2]. Life-history parameters (LHP), such as pregnancies, skeletal trauma, and renal disease, can be identified and precisely datable from incremental lines of AEFC in human teeth by observing their visual effects [3]. These life history parameters appear to change calcium metabolism [4, 5], and lack of available calcium at the mineralization front of the cementum causes formation of a such visually different incremental AEFC line [3]. In a study on humans [3], as well as in great apes [6] “suspicious” AEFC lines were successfully detected as being visibly broader and translucent in tooth ground sections (70 - 80μm thick) under optical magnification with transmuted polarized light. These studies also showed that some of the LHPs affecting mineralization of AEFC are precisely datable from the AEFC cross-sections. On the other hand, in a controlled study undertaken on goats, Lieberman [2] showed that AEFC bands corresponding to a control diet low in minerals including calcium and phosphorus appeared to be opaquer and relatively narrower, as observed from x-ray microradiographs of thin ground sections (50μm thick). Lieberman described these bands as hypermineralized (denser) due to reduced cementogeniesis. In contrast, a study undertaken by Cool and colleagues [7] reported that cementum growth layers are not the result of changes in mineral density at all, as they failed to detect cementum growth layers using scanning electron microscope (SEM) equipped with backscattered electrons detector (BSE). The most recent study on composition and structure of AEFC, using Raman imaging analysis [8] argues that darker AEFC lines correspond to higher mineral/organic ratio when compared to brighter lines.

However, due to the relatively regular annual rhythm in their layering, AEFC incremental lines are more frequently used as a chronological age estimation aid. This optical detection technique has been in relatively frequent use as an individual age estimation aid [9 – 21], although many results are carefully qualified or subsequently disputed [22, 23]. Continued caution is required, as cementum is the least known of all the mineralized tissues [1], and rigorously controlled human clinical studies have rarely been used to support these findings. As such, life history researchers cannot entirely rely on the results of these pioneering studies yet. Furthermore, cementum research is considerably hampered by an over-emphasis on optical microscopy as summarised by Nadji and colleagues [24]. We do not fully understand the optical appearance of cementum incremental lines yet, let alone the underlying complex mineralisation process(es).

Here we investigate if chemical composition and the degree of mineralization of AEFC can detect one important LHP, namely full-term pregnancies, from human teeth. To do so, we employ a comparative approach towards the study of AEFC incremental lines. Firstly, we compared direct measurements of degree and distribution of mineralization of AEFC from a patient with a known life history of six full term pregnancies, using two different microscopic methods, Scanning Electron Microscopy (SEM) with energy Dispersive X-Ray Analysis (EDS) and Time-of-Flight Secondary Ion Mass Spectroscopy (ToF-SIMS).

## Material and methods

A mandibular canine was extracted from a white Caucasian woman undergoing necessary dental intervention at the University Hospital Kosovska Mitrovica (University of Priština). The informed consent to use the tooth for this research was obtained from the patient as well as her anamnestic data (Table 1). The patient was born and raised in the region of Kosovska Mitrovica. At the time of the extraction she was 66 years old with no previous history of renal disease, endocrinal problems, skeletal fractures or trauma. The patient was considered an excellent candidate for a fertility related analysis, as she reported six pregnancies that carried to full-term, starting at the age of 19 with the last one at age of 31 (Table 1). After the extraction, the tooth was placed in a labelled vial containing physiological saline (solution of 0.90% w/v of NaCl). The tooth was free from obvious signs of pathology.

**Table 1.**
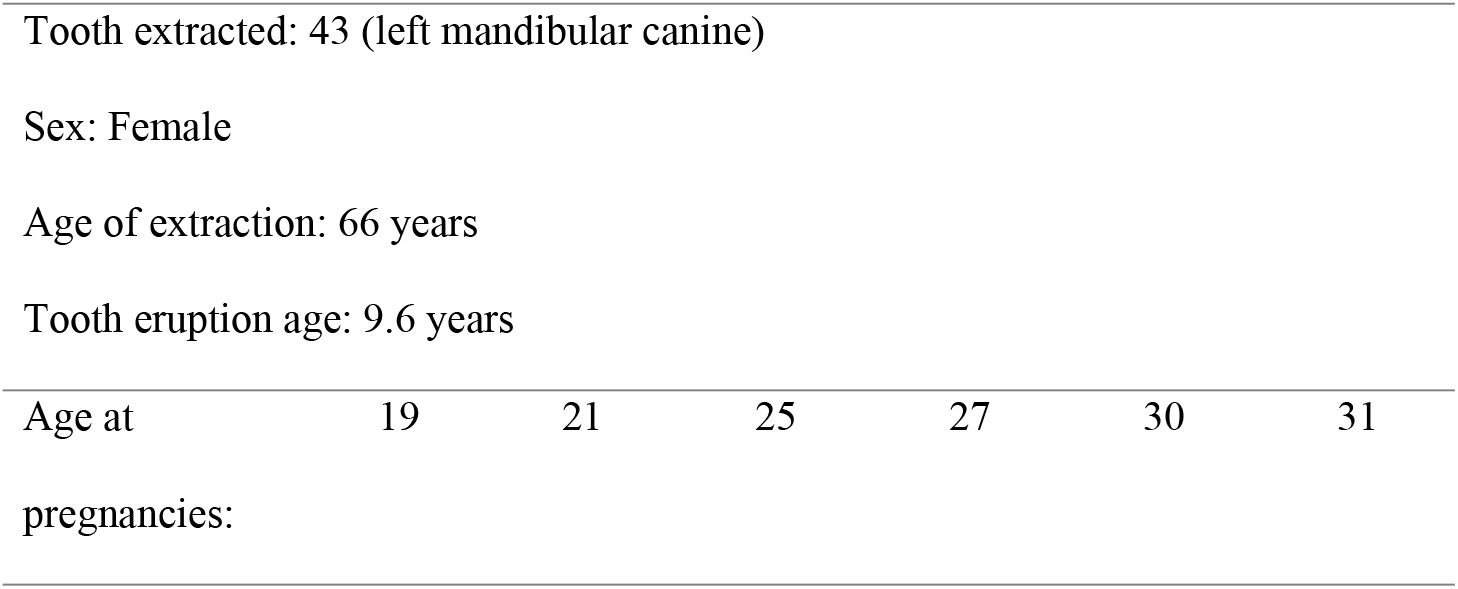
The patient’s anamnestic data

### Scanning Electron Microscopy (SEM) with Energy Dispersive X-Ray Analysis (EDS) for measurement of elements

The resin block with exposed mid-root surface represented the sample to be analysed by SEM-EDS was coated with carbon (Quorum K975x carbon coater; Quorum Technologies, UK). The prepared sample was examined at 20 keV by scanning electron microscopy using a Philips XL30 E-SEM (Hillsboro, OR, USA) equipped with an Oxford instruments energy dispersive x-ray analysis detector using Oxford Instruments INCA software. The EDS analysis was employed to determine whether there are mineral component compositional changes between cementum growth layers. EDS analysis was performed on the same sections employed for SIMS imaging. Beam positioning was achieved by viewing the BSE image at 500× magnification.

Line scans of cementum growth layers was subject to an acquisition time of 100 sec (working distance 10 mm, take-off angle 35°) to obtain X-ray spectra. The X-ray spectra were used to determine which minerals were present and the Ca:P atomic percent ratio.

### Identification of Ca and HAp by Time of Flight–Secondary Ion Mass Spectrometry Imaging

The sample preparation for ToF-SIMS imaging comprised several steps and it was performed at University College London, Institute of Archaeology. The procedure was tailor made for this research, namely the identification of Ca and HAp form AEFC growth layers using ToF-SIMS. After the cross section was cut out from the mid-root of the patient’s tooth, the exposed root surface mounted in the resin represented the sample to be analysed using ToF-SIMS. The following preparation step was polishing, undertaken using a rotating wheel and polishing media. This step is required to remove the surface damage that occurred during sectioning and to provide a flat surface. The polishing procedure included the use of a series of progressively finer polishing pads and diamond compounds, from 2 − ¼ μ (Kemet Diamond paste). The final step was ultrasonic cleaning (1min at xx kHz) using deionized water. This step was performed in order to thoroughly remove all traces of contamination tightly adhering or embedded onto the sample surface.

Secondary ion mass spectrometry and secondary ion mapping was performed using a TOF.SIMS5 mass spectrometer (ION-TOF, Münster, Germany) at Imperial College London, Micrometre resolution was used for secondary ion mapping within m/z 0-880. The system is comprised of a bismuth primary ion beam, operating at 25 keV and tuned to use the Bi_3_ ^+^ cluster for greater secondary ion yield, and a low energy electron flood gun for charge compensation. Cluster ion sources, such as Bi_3_, are used to identify larger HAp fragment ions at for example m/z 485, 541, 597, and 653, identified as Ca_5_P_3_O_12_^+^, Ca_6_P_3_O_13_^+^, Ca_7_P_3_O_14_^+^, and Ca_8_P_3_O_15_^+^, respectively. Ionic species sputtered from the surface under the bismuth bombardment are steered into a reflectron time-of-flight mass analyzer. Before mass spectrometry was performed, an Ar_n_^+^ cluster ion beam was used to remove any surface organic contaminants. Identified peaks strongly localized to cementum growth layers were mapped on single ion maps. Positive-ion spectra were acquired from two different 100×100 μm regions of tooth encompassing the entire cementum width from mesio-buccal and disto-buccal side of the tooth, respectively to localize of HAp and identification of different CaP phases within cementum layers.

### Study Approval

Use of human tissues and human sensitive data for this study was approved by the Ethics Committee of Faculty of Medicine, University of Pristina at Kosovska Mitrovica, Ministry of Health, Republic of Serbia, as well as by the Ethical Board of Research Executive Agency, European Commission, Brussels.

## Results

### SEM-EDS

No visual evidence of cementum growth layering was found SEM micrograph using Philips XL30 FEG-SEM (Hillsboro, OR, USA) equipped with an Oxford instruments energy dispersive x-ray analysis detector (Fig 1a). The Ca:P ratio (by atomic percent) ranged from 1.47 to 1.73, with 1.59 as average value. Line scan for Ca showed no significant change in its relative amounts across the width of the cementum (Fig 1b), except in the case of line spectrum (8) which exhibits the lowest relative amount of Ca as of 0.13(atomic %), as well as the lowest Ca:Pa ratio as of 1.29 by atomic % (Fig.1a). However, these low readings for the line spectrum (8) are due to an intruding artefact deposited in cementum, which can be clearly observed on electron photomicrograph of a transverse section of midroot cementum (Fig 1a), and should not be taken into account when interpreting mineral distribution across the AEFC width of our sample.

**Fig 1.**
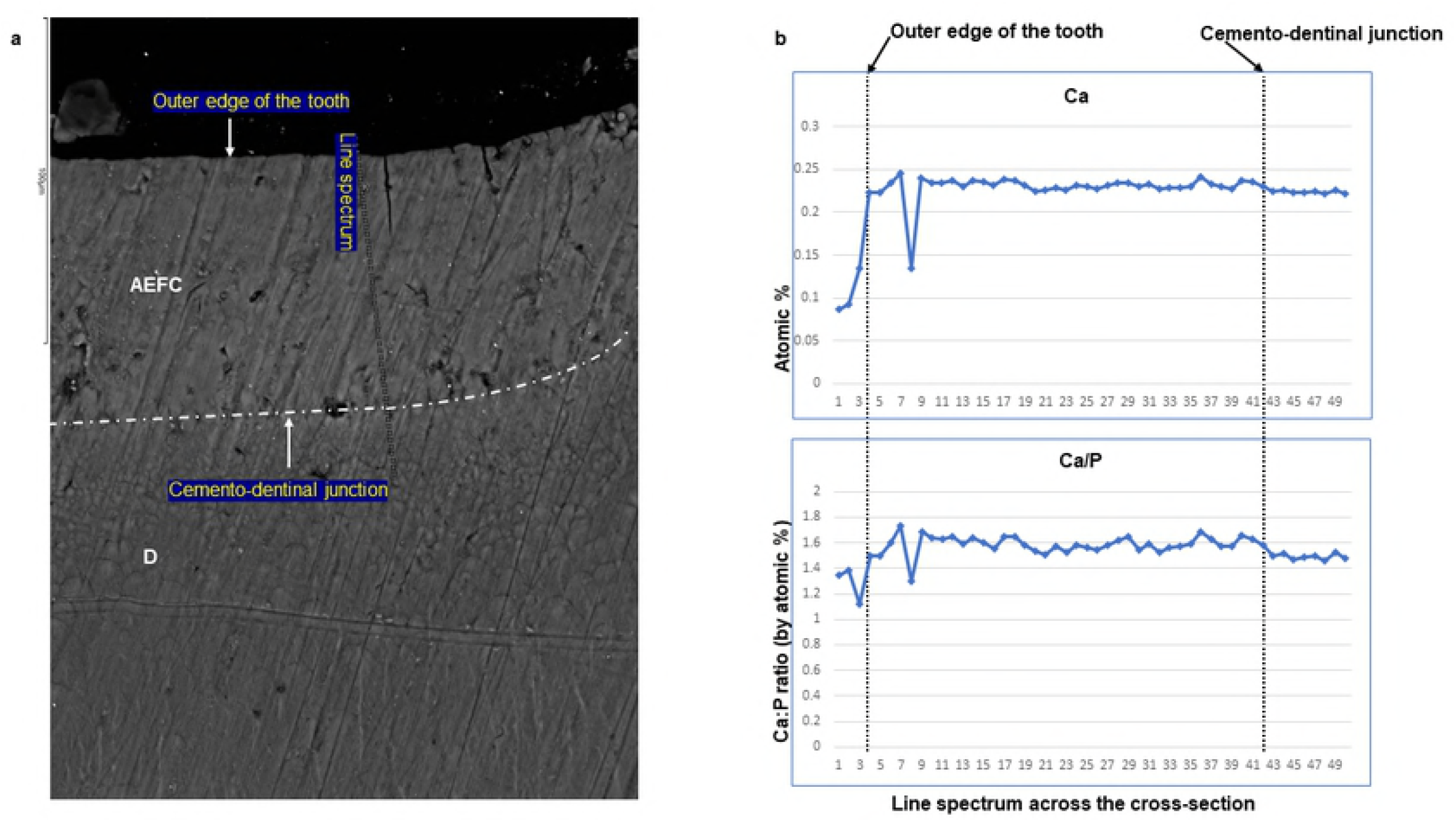
Results of EDS line analysis of the cross section of the patient’s tooth. The SEM photomicrograph of the cross section (a) is showing the location across mid-root acellular extrinsic fiber cementum (AEFC) and dentine (D) where the line spectrum was taken. The line charts (b) represent results of EDS line scan analyses for Ca (upper chart) and Ca/P ratio (lower chart) across the width of AEFC.

### ToF-SIMS

Layering of AEFC is somewhat visible from the ION-TOF.SIMS5 camera view (Fig 2a), but not in a form of clearly defined incremental lines. Furthermore, AEFC incremental lines in our sample were not clearly discernible even when observed under transmitted polarized light microscope, using 400× magnification, and flowing the established Tooth Cementum Annulation protocol [25] (see S1 Appendix and S1 Fig). However, we were able to estimate approximal width of AEFC incremental lines in our sample by using the available measurements, and knowledge on teeth eruption AEFC annulation. Having the total width of the intact cementum layer measured from the ToF-SIMS micrograph (Fig 2a) of our sample, assuming that it grows in a regular annual rhythm [1 – 3], we were able to calculate the approximate width of incremental lines. As shown on Figure 2a, the analysed AEFC is flanked by cemento-dentinal junction on one side and outer edge of the tooth on the other side. More precisely, the total width of AEFC in our sample equals 73μm, as it spreads across the area between 4 μm - 77 μm on the linescans (Fig 2b). Given the age of extraction for this tooth (66 years), as well as the average sex specific year of eruption of the tooth, which is 9.6 for mandibular canines in females [26, 27], the estimated approximal width of AEFC incremental lines for this individual is c. 1.3 μm.

**Fig 2.**
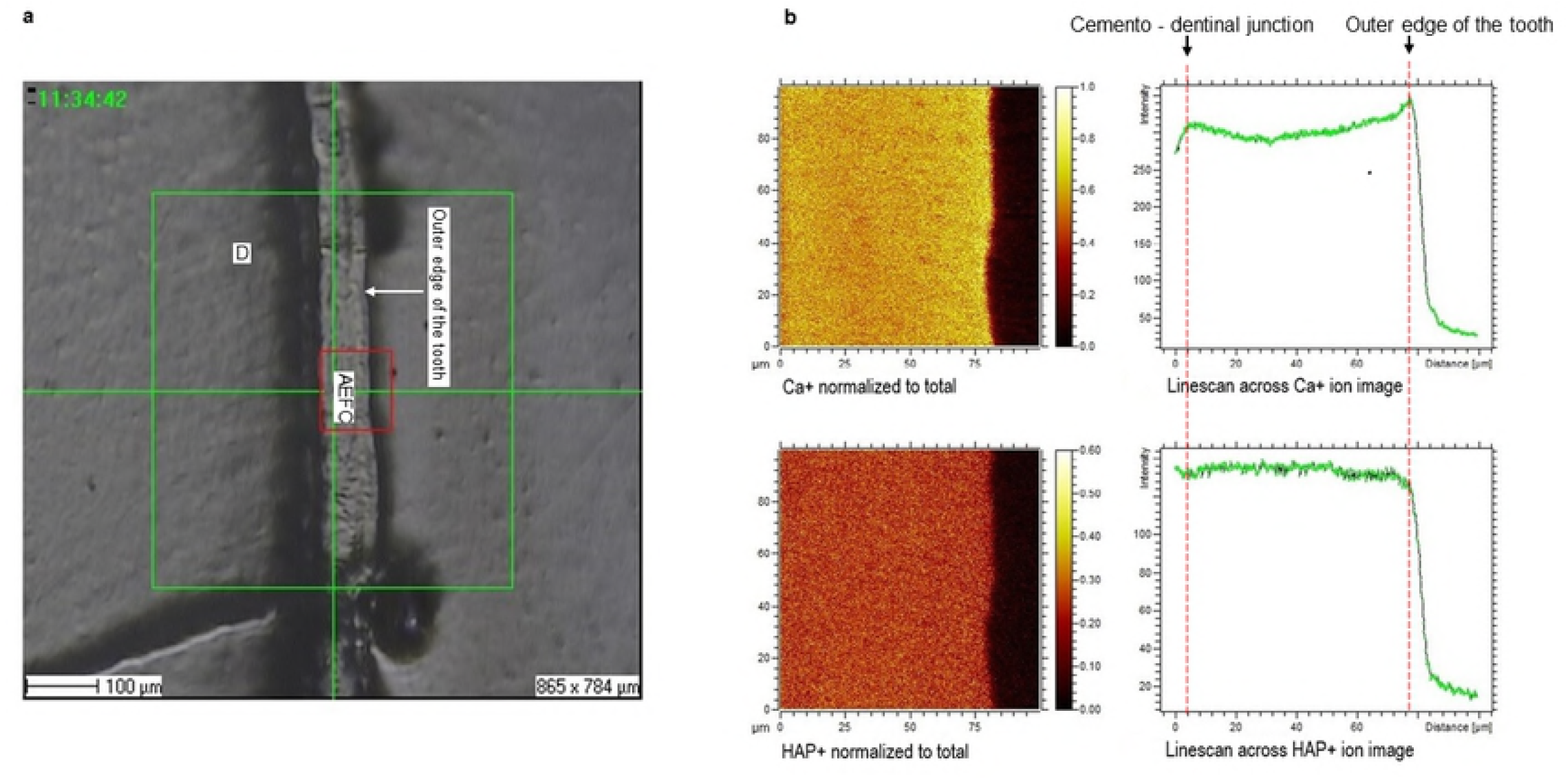
Distribution of the molecular ions identified by SIMS. (a) A ToF-SIMS micrograph of the area of interest. “D” represents dentine; “AEFC” represents acellular extrinsic fiber cementum; and red square is demarking the area which has been analysed by ToF-SIMS. (b) Molecular ion mapping and line scan ion images of calcium (Ca+), and (C) hydroxyapatite (HAp+).

Elemental and molecular maps, as well as line scans of Ca+ and HAp+ (Ca/P ratio) are obtained from the AEFC surface. We have detected a variation in the intensity of Ca+ across the analyzed AEFC surface (Fig 2b). A depletion in relative Ca+ intensity can be observed from 12 – 54μm respectively, but no depletion in intensity of HAp+ in the same line scan (Fig 2b, Fig 3). This corresponds to the patient’s 2^nd^ -5^th^ decade of life (Fig 3). The lowest point in Ca+ intensity depletion is recorded at 32^nd^ μm (Fig 3) which corresponds to start of the patent’s 4^th^ decade of life (around age her age of 30).

**Fig 3.**
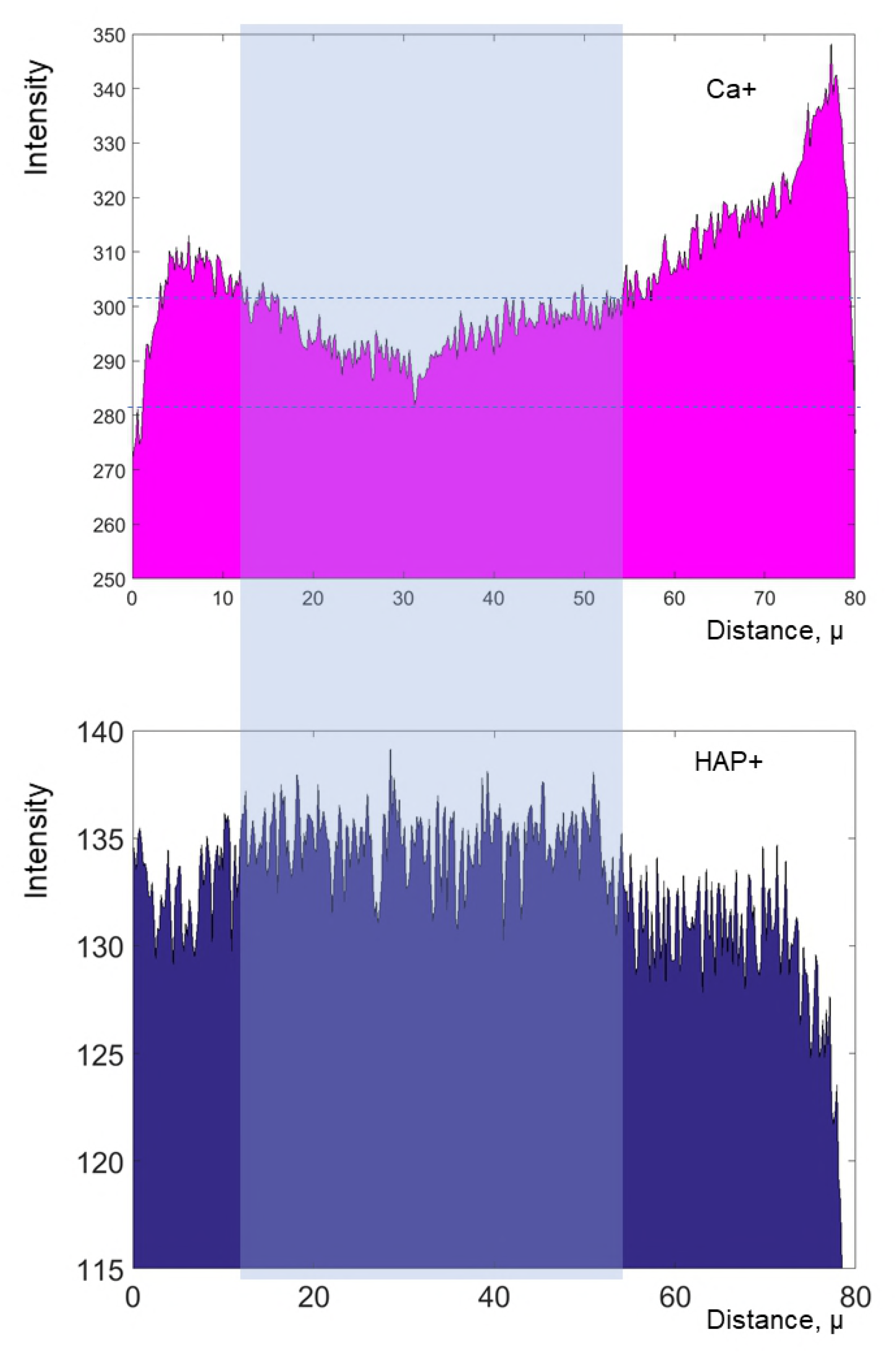
Secondary ion yield (counts) of calcium (Ca+) and hydroxyapatite (HAp+) across the width of the cementum. Right side of the plots represents outside of the tooth.

## Discussion and conclusions

This study compared two different elemental analyses methods in order to establish which one best estimates the degree and distribution of mineralisation of AEFC. The results of the two methods were compared with the recorded life history parameters of a subject who had six full-term pregnancies. We demonstrate that degree and distribution of mineralization of human AEFC varies across the width of the resultant cementum sample. Furthermore, we show how life history parameter detection in AEFC can vary between two elemental detection methods.

Scanning electron microscope with electro-dispersive probe did not detect any significant variation in Ca relative amounts across the AEFC width, nor in Ca:P ratio respectively (Fig 1b). The range for the Ca/P ratio (by atomic percent) varied between 1.47 to 1.73, where the majority of the values (Fig 1b) fell below 1.65, and the value 1.73 was read only once. This implies that our results for Ca/P atomic percent ratio are significantly lower than the Ca/P atomic ratio bioapatite standard (1.69 – 1.71). These results suggest that the AEFC analysed here is relatively hypomineralized overall. The aim of this study was to detect variation in degree and distribution of mineralization across the AEFC cross-section, therefore, it is not of any use to discuss the average Ca/P ratio we have obtained. In terms of previous research that found no obvious variation in the concentration profile for calcium and phosphorus, our results are in general agreement [7]. The authors of that study reported that calcium and phosphorus are present in cementum at a ratio (1.70) similar to the bioapatite standard, which disagrees with our results. They concluded that “cementum growth involves a constant rate of both mineralization and matrix production, rather than variations in the rate of matrix production with mineralization continuing at a uniform rate” [7], which is opposite of what was reported in Lieberamn’s work [28] (Lieberman, 1994). Apart from stating that the cross-section was taken through areas shown to contain growth layers, the authors [7] had not specified precisely which type of cementum analyzed in their study. This implies that they were observing cementum as a single uniform type of tissue. A similar approach was taken in a few previous studies when analyzing cementum with electron probes [29, 30]. Importantly, there are various types of cementum which can be distinctly classified based on presence or absence of cells, nature and origin of organic matrix, or a combination of these factors [1, 31, 32]. Different types of cementum are formed at different rates, but only AEFC is considered annually deposited. Most importantly, incremental lines are found in different types of cementum (e.g. in acellular and cellular cementum). Furthermore, EDS analysis can only be used to obtain the relative amounts of concretions even when certified standards are used; as EDS operating software does not provide us with continuous elemental values. This might be the reason we were not able to detect any obvious variation in Ca and P profiles. With all this in mind, we argue that the SEM-EDS technique is inadequate for measuring the degree and distribution of AEFC mineralization.

ToF-SIMS micro-image revealed clearly defined AEFC in our cross-section. Although some appearance of layering within AEFC can be also observed (Fig 2a), the incremental lines of AEFC were not clearly discernible from the micrograph. On the other hand, we were able to estimate the approximate width of AEFC incremental lines using the micrograph itself. The linescans showed obvious variation in Ca+ intensity across the AEFC width, but not for HAp+ (Fig 2b, Fig 3). Ca+ intensity variation has been detected in a form of a depletion which might correlate with the with patient’s pregnancies, as the depletion of Ca+ intensity corresponds to the patient’s 2^nd^ – 5^th^ life decade (Fig 3). The lowest point in Ca+ intensity depletion is recorded at 32^nd^ μm which corresponds to the beginning of the patent’s 4^th^ life decade. The initial Ca+ intensity depletion was detected four years prior to the patient’s age at the time of the first pregnancy. The lowest value of Ca intensity is measured around the age corresponding to the patient’s last pregnancy. From that point on, the intensity values for Ca+ are seen to rise or perhaps normalise, after final pregnancy.

In conclusion, our ToF-SIMS results imply that life history parameters, such as pregnancies, are more likely to influence AEFC in terms of relatively reduced mineralization, which is in accordance with work done by Kagerer and Grupe [3]. We also demonstrated how use of SEM-EDS technique is inadequate when measuring the degree and distribution of AEFC mineralization. Our results point towards an alternative direction for life history research using tooth cementum data. To detect life history parameters, which have a marked impact on Ca metabolism, such as pregnancies, we propose a new methodological approach, namely Time-of-Flight Secondary Ion Mass Spectrometry. As a clear relationship between degree and distribution of AEFC mineralization and reported pregnancies is observed by our study, this technique appears to have great potential for investigating various biological events in the historical development of humans and animals. Clearly, more clinical tests are required. In the meantime, far more caution is required by cementum-oriented life history researchers.

## Supporting Information

**S1 Appendix. Transmitted Polarized Light Microscopy Analysis**

**S1 Table. Results of the reported counts of incremental lines from the tooth cross-section.**

**S1 Fig. Ground cross-sections of the patient’s tooth under the transmitted polarized light microscope (a and b ×400).** The red arrows indicate a pronounced eruption line. The white arrows indicate possible “crisis” lines which appear as broad and translucent layers in acellular extrinsic fiber cementum (AEFC) of the patient. (D) represents dentine, and (CDJ) is cemento-dentinal junction.

